# Researchers collaborate with same-gendered colleagues more often than expected across the life sciences

**DOI:** 10.1101/345975

**Authors:** Luke Holman, Claire Morandin

## Abstract

Evidence suggests that women in academia are hindered by conscious and unconscious biases, and often feel excluded from formal and informal opportunities for research collaboration. In addition to ensuring fairness and helping to redress gender imbalance in the academic workforce, increasing women’s access to collaboration could help scientific progress by drawing on more of the available human capital. Here, we test whether researchers tend to collaborate with same-gendered colleagues, using more stringent methods and a larger dataset than in past work. Our results reaffirm that researchers co-publish with colleagues of the same gender more often than expected by chance, and show that this ‘gender homophily’ is slightly stronger today than it was 10 years ago. Contrary to our expectations, we found no evidence that homophily is driven mostly by senior academics, and no evidence that homophily is stronger in fields where women are in the minority. Interestingly, journals with a high impact factor for their discipline tended to have comparatively low homophily, as predicted if mixed-gender teams produce better research. We discuss some potential causes of gender homophily in science.

## Introduction

Women are severely underrepresented in many branches of science, technology, engineering, mathematics, and medicine (STEMM), and face additional challenges and inequities relative to men [1–5]. On average, women occupy more junior positions [6,7] with lower salaries [8,9], receive less grant money [10,11], are promoted more slowly [12–15], and are allocated fewer resources [16] and less research funding [17–19]. Experimental evidence suggests that bias against women plays a major role in generating these differences [20,21].

Writing papers, networking, and collaboration are all instrumental to research productivity and academic career advancement [22–25], and dozens of studies have tested for gender differences in these areas [5,26–29]. For example, studies have concluded that women tend to be less involved in international collaboration [19,28,30–32], collaborate less within their own university departments [31], have less prestigious collaborations [33], and fewer collaborations in total [34]. These gender differences in collaboration practice presumably have multiple causes, which might include implicit and explicit gender bias [20], differential family obligations [33,35,36], gender differences in confidence or self-esteem [37], concerns relating to sexual harassment [38], and unequal access to conferences [39] or travel funds [32].

A high, steadily increasing proportion of research papers is written by more than one author [3], making collaboration a key predictor of publication output, and thus of career prospects [40,41]. Additionally, empirical studies imply that mixed-gender or otherwise diverse teams produce better outputs on collaborative tasks than less diverse teams [42–48]. For reasons such as these, multiple studies have examined the author lists of published research articles in order to test for gender differences in collaboration frequency or pattern. To our knowledge, most or all such studies imply that men co-publish with men, and women with women, more often than expected if collaborators assort randomly with respect to gender [49–58]. This pattern of assortative publishing has often been termed ‘gender homophily’.

However, we believe that earlier studies of gender homophily were hindered by a largely unacknowledged statistical issue that we will refer to as the Wahlund effect (Figure 1), by analogy with the conceptually similar Whalund effect in population genetics [59]. The Wahlund effect makes it deceptively difficult to infer gender-based preferences simply by counting the number of same‐ and mixed-gender coauthorships. Essentially, whenever coauthorship data are sampled from two or more discrete sets of literature, which vary in the author gender ratio and which are largely not connected by collaboration, the number of same-gendered coauthors will be inflated. This can give the impression that authors preferentially publish with same-gendered colleagues even if no gender preferences exist, or if the true preference is for opposite-gendered colleagues (‘gender heterophily’). For example, a sample of literature containing a mixture of bioinformatics and cell biology papers will probably contain an excess of mostly-male and mostly-female author lists, simply because researchers usually collaborate within their own discipline, and because the author gender ratio is more male-biased in bioinformatics than in cell biology [5].

**Figure 1:**
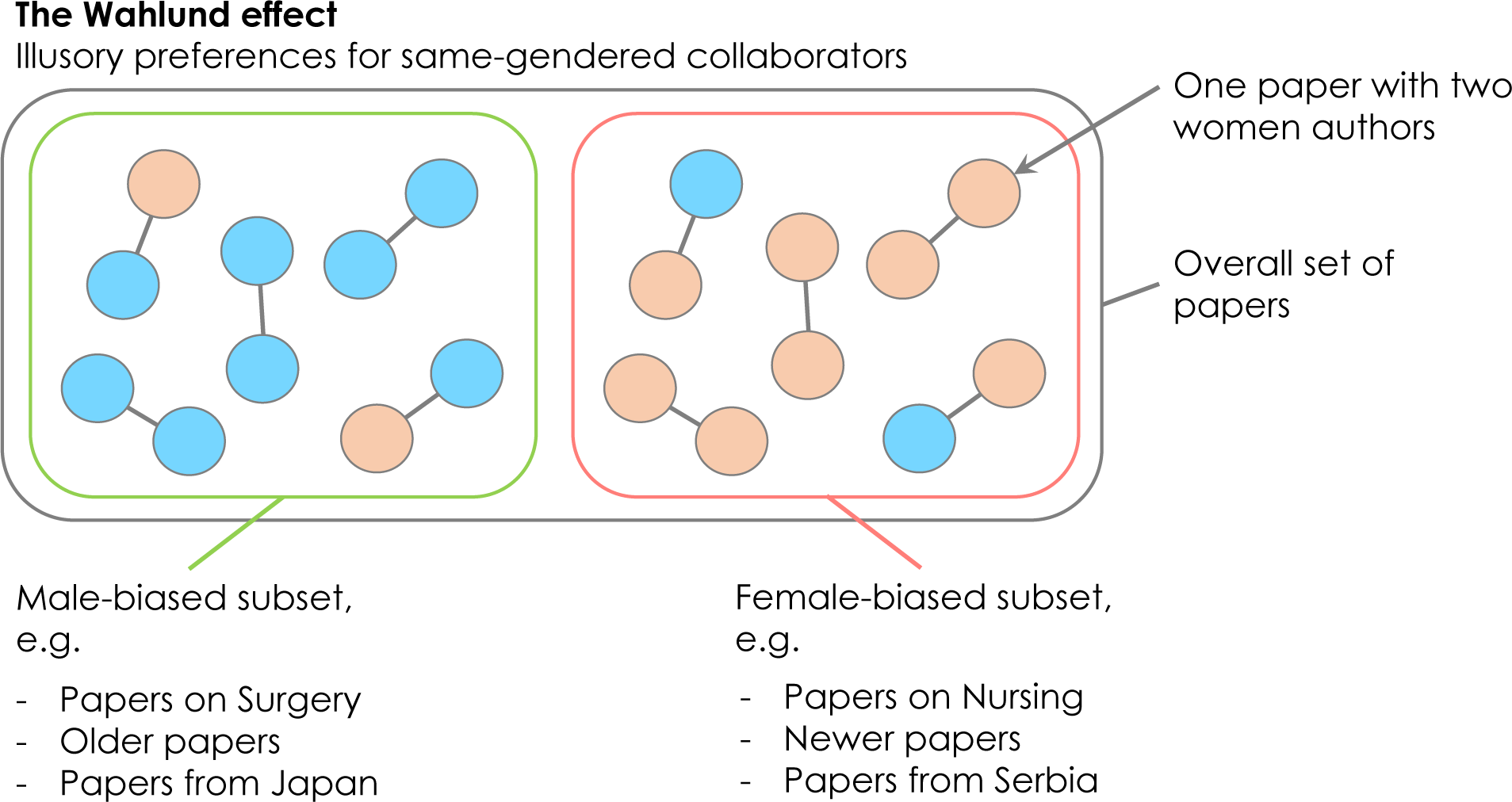
The Wahlund effect can make it appear as if authors publish with same-gendered colleagues disproportionately often, even if collaboration is completely random with respect to gender. Here, coloured circles represent male and female authors, and coauthors are linked with lines. Across the whole set of ten papers, there is an apparent excess of same-gender collaborations: there are six same-gender papers and only four mixed-gender papers, which is fewer than the 10 *×* 2 *×* 0.5 *×* 0.5 = 5 mixed-gender papers expected under the null hypothesis that authors assort randomly. However, within each subset, there is no evidence that authors prefer to publish with same-gendered individuals (if anything, this small dataset suggests gender heterophily). The Wahlund effect will tend to inflate the frequency of same-gender coauthorships whenever the data is composed of two or more disconnected subsets of literature with different author gender ratios; these subsets could be research disciplines, older versus newer papers, or papers from authors in different countries. The example countries and disciplines were selected based on data in [5].

In the present study, we test whether life sciences researchers tend to co-publish with same-gendered colleagues, while controlling for the Wahlund effect as strictly as possible. We use a recently-published dataset describing the gender of 35.5 million authors from 9.15 million articles indexed on PubMed [5]. Holman et al. [5] reported large differences in the gender ratio of authors across research disciplines, journals, countries, and across the years 2002-2016. We therefore tested for gender homophily while restricting our analysis to particular journals (i.e. research specialties), time periods, and countries. We quantified gender assortment using a metric called *α′* [60], which is positive when same-gender authors publish together more often than expected (gender homophily), negative when opposite-gender authors publish together more often than expected (heterophily), and equal to zero when authors assort randomly with respect to gender (see Methods).

## Results

### Gender homophily by discipline, time period, and authorship position

Figure 2 shows the distribution of *α′* estimates in 2015-2016 across all journals for which we recovered sufficient data, when *α′* was calculated for all authors, first authors only, or last authors only. Most journals had positive values of *α′* (77-92%, depending on time period and author type; S1 Data), and for many of these the false discovery rate (FDR)-corrected p-values suggested that *α′* was significantly greater than zero (1469/2077 journals were significant in 2015-16, and 404/1192 in 2005-6; S1 Data). Only 2/2077 journals had statistically significant heterophily (i.e. *α′* < 0) in 2015-16, and 1/1192 in 2005-6 (S2 Table). The remaining 606 or 787 journals (in 2015 and 2005 respectively) had a value of *α′* not significantly different from zero, consistent with the null hypothesis of random assortment with respect to gender. We also confirmed that in most journals (S2 Data) and most research disciplines (S3 Data, S1 Fig), the majority of papers had multiple authors.

**Figure 2:**
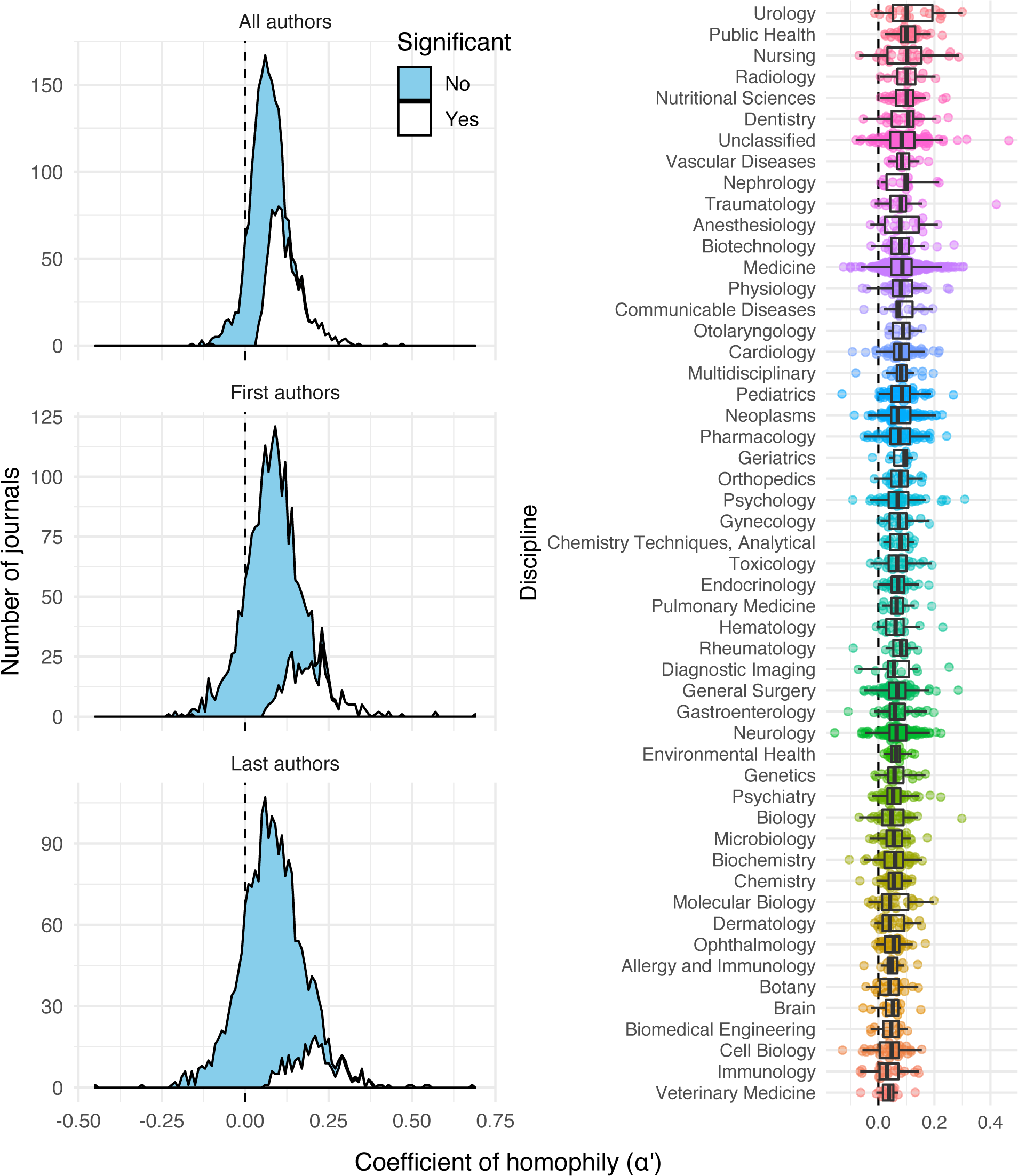
Of the 2116 journals for which we had adequate data in 2015-2016, 825 showed statistically significant evidence of gender homophily (denoted by *α′* > 0), and 1 showed statistically significant evidence of heterophily (*α′* < 0), after false discovery rate correction. In the stacked density plot, the white area shows the number of journals for which homophily was significantly stronger than expected under the null hypothesis (corrected p < 0.05), while the blue area shows all the remainder. Patterns were similar whether *α′* was calculated for all authors, for first authors only, or for last authors only. Points in the right panel show *α′* for individual journals.

*α′* was significantly higher in the literature sample from 2015-16 relative to 2005-6, though the difference in means was small (S2 Fig; Effect of the fixed factor ‘Time period’ in a linear mixed model of the data for all author positions: Cohen’s *d* = 0.091*±*0.04, *t*_953_ = 2.42, p = 0.016).

When comparing pairs of *α′* values estimated for the first and last authors for the same journals, we found that *α′* tended to be higher for first authors than for last authors (S3 Fig; Effect of the fixed factor ‘Authorship position’ in a linear mixed model: Cohen’s *d* = 0.065±0.02, *t*_2024_ = 4.28, p < 0.0001). This suggests that the gender of the first author was a slightly stronger predictor of the remaining authors’ genders than the gender of the last author, i.e. the opposite of what is predicted if senior scientists are causally responsible for homophily.

### Variance in homophily between disciplines

Figure 2 illustrates the variance in journal homophily values (*α′*) across scientific disciplines. All disciplines had positive mean *α′* (averaged over journals), although homophily appeared somewhat stronger in some disciplines than others (e.g. mean *α′* was 0.12*±*0.02 for Urology journals and 0.03*±*0.01 for Veterinary Medicine journals; Figure 2, S4 Data). However, there was no formal evidence for consistent differences in *α′* between disciplines: the random factor ‘Discipline’ explained around 1% of the variance in *α′* in the two linear mixed models described in the previous section (see Figure 2 and mixed models in Online Supplementary Material). Thus, whatever processes are responsible for producing positive *α′* values appear to be similarly strong in all the disciplines we examined.

There was no indication that journals publishing on a wide range of topics have higher *α′* values than more specialised journals, due to the Wahlund effect. For example, the journal category ‘Multidisciplinary’ – which includes journals like *PLOS ONE*, *Nature*, *Science*, and *PNAS* – did not have markedly elevated *α′* (Figure 2). This result suggests that our estimates of homophily, and estimates from some of the earlier studies listed in the Introduction, are probably not inflated by the presence of disparate research topics (with variable author gender ratios) being published within individual journals.

Nevertheless, when we calculated *α* across all non-single-author papers in our entire 15-year PubMed dataset (as before, excluding papers where at least one author’s gender was unknown; n = >3 million papers, >16 million authors), we found that *α* was 0.126. This figure is almost double the median value of *α′* for individual journals (Figure 2; *α′* = 0.070 for ‘All authors’), suggesting that lumping together papers from different fields and different time periods can indeed produce spurious evidence for gender homophily as outlined in Figure 1.

### Relationship between gender homophily and number of authors

Papers with two authors had significantly lower (but still positive) *α′* values relative to papers with more than two authors, on average (Figure 3; statistical results in Online Supplementary Material). Papers with 3, 4 or *≥* 5 authors had essentially identical average *α′* values. The variance in *α′* across journals was also a little higher for 2-authors papers compared to the remainder (Figure 3), though part of this variance is due to the reduced sample size (in terms of number of authors) for the 2-author papers. One possible explanation for this finding is that 2-authors papers might be more likely to have an author list that is evenly split between career stages (e.g. a postgraduate student and their supervisor), increasing the chance that the authors are mixed gender (see Figure 6). The result also suggests that the processes responsible for gender homophily are similar in small (e.g. 3 author) and larger (5+ authors) collaborations (and across disciplines where small versus large collaborations are the norm).

**Figure 3:**
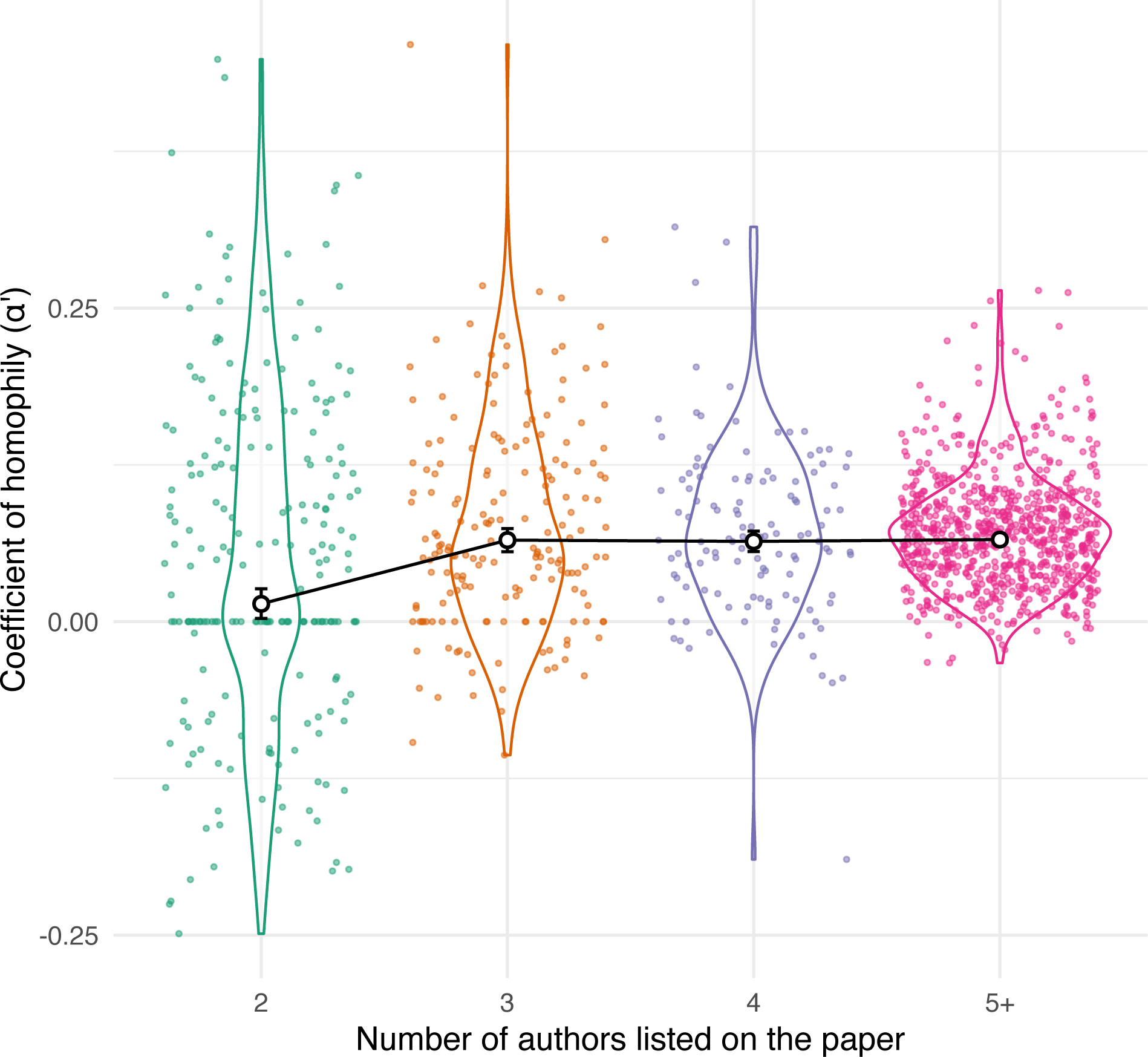
The coefficient of homophily (*α′*) was slightly less positive when calculated for two-author papers only, relative to papers with longer author lists. The individual points, whose distribution is summarised by the violin plots, correspond to individual journals. The larger white points show the mean for each group (and its 95% CIs), as calculated by a Bayesian meta-regression model accounting for repeated measures of *α′* within journals, as well as the precision with which *α′* was estimated.

### Relationship between gender homophily and gender ratio

We next tested whether researchers are more or less likely to publish with same-gendered colleagues in strongly gender-biased disciplines (e.g. Surgery or Nursing), relative to disciplines with a comparatively gender-balanced workforce (e.g. Psychiatry). We found a positive, non-linear relationship between the overall gender ratio of all authors publishing in a particular journal [5], and the estimated value of *α′* for all authors and for first authors (Figure 4). Journals with a balanced or female-biased author gender ratio tended to have higher *α′* (i.e. stronger homophily) than journals with a male-biased author gender ratio (GAM smooth terms p < 0.001; Online Supplementary Material). The relationship was not statistically significant when *α′* was calculated for last authors (GAM, p = 0.142), though the trend appeared similar (Figure 4).

**Figure 4:**
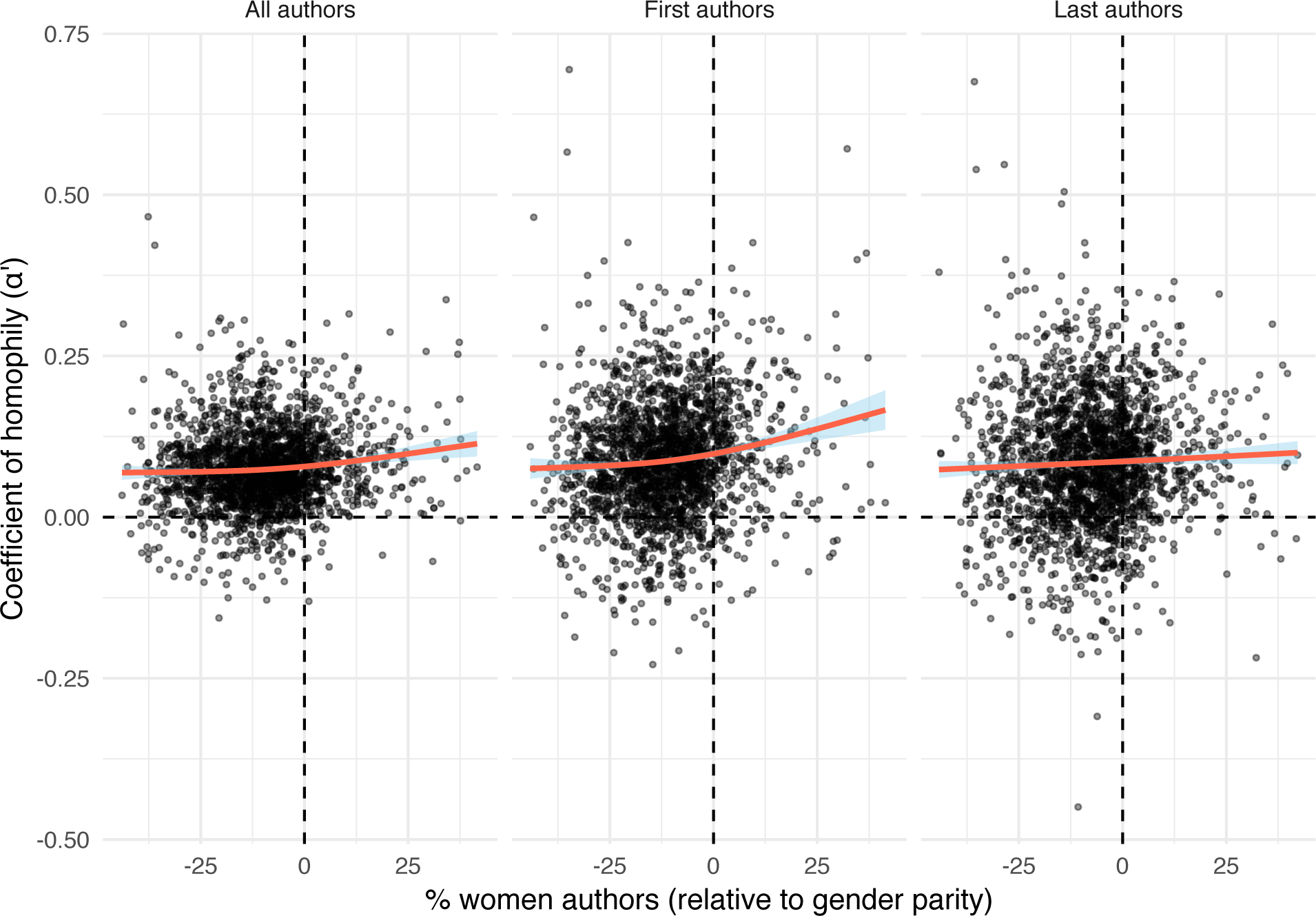
There is a weakly positive, non-linear relationship between the gender ratio of authors publishing in a journal, and the coefficient of homophily (*α′*). Specifically, journals with 50% women authors or higher tended to have more same-sex coauthorships than did journals with predominantly men authors. This relationship held whether *α′* was calculated for all authors, first authors only, or last authors only. A negative value on the x-axis denotes an excess of men authors, a positive value denotes an excess of women authors, and zero denotes gender parity (i.e. equal numbers of men and women). The lines were fitted using generalised additive models with the smoothing parameter *k* set to 3.

### Relationship between journal impact factor and gender homophily

We observed a noisy but statistically significant linear relationship between standardised journal impact factor and *α′*, such that journals with a high impact factor for their discipline had weaker gender homophily than did journals with a low impact factor for their discipline (Figure 5; linear regression: *R*^2^ = 0.043, *t*_1415_ = −8.0, p < 0.0001). The slope of the regression was *−*0.012*±*0.0015, indicating that increasing the discipline-standardised impact factor by one standard deviation is associated with a reduction in *α′* of 0.012.

**Figure 5:**
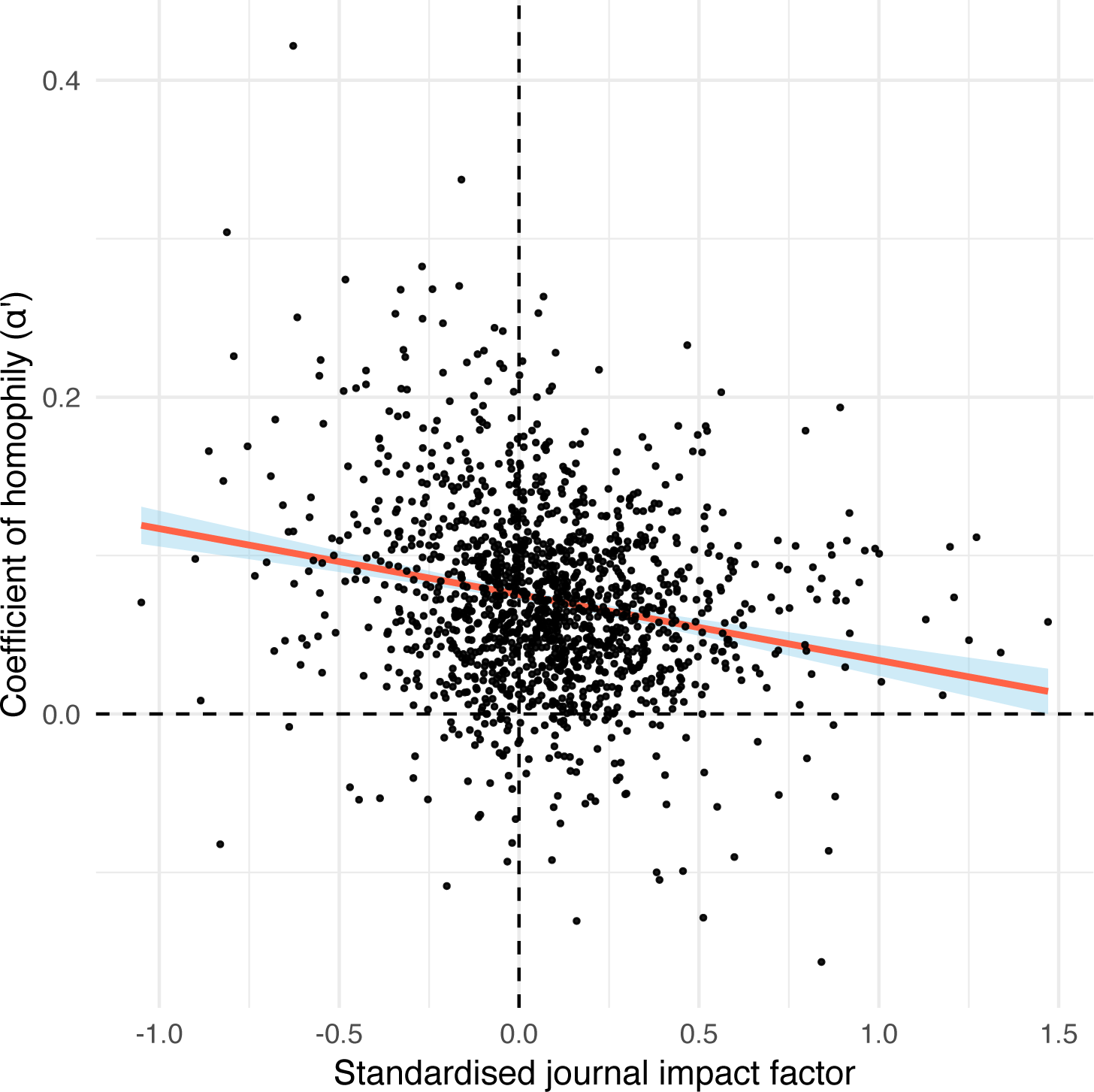
Journal impact factor (expressed relative to the average for the discipline) is negatively correlated with *α′*. The relationship is noisy (*R*^2^ = 0.043), but the results suggest that journals with strong homophily tend to have lower impact factors than journals with weak homophily in the same discipline.

### Analysis accounting for differences in author gender ratio between countries

When we restricted the analysis by country, we observed statistically significant homophily for 72 of the 325 journal-country combinations tested (64 unique journals and 18 unique countries), and no significant heterophily (S4-S5 Fig). Additionally, the values of *α′* calculated for each journal-country combination were only very slightly lower than the *α′* values calculated for the journal as a whole (i.e. when pooling papers from different countries, as was done to make Figure 2): on average, the difference in *α′* was only 0.002 (S6 Fig). These results suggest that our findings of widespread homophily in the main analysis were not driven solely by a Wahlund effect resulting from gender differences between countries.

### Theoretical expectations for *α* when the gender ratio differs between career stages

Given that we cannot identify individual researchers or their career stages, we used a simple model to derive the theoretical expectations for *α* when the gender ratio differs between career stages (see Methods). As shown in Figure 6, we predict that *α* is expected to be non-zero, even if collaborators are randomly selected with respect to gender, provided that there is a gender gap between career stages. The extent to which *α* deviates from zero depends on the relative frequencies of collaboration within and between career stages (rows and columns in Figure 6), and the size of the gender gap between stages (x‐ and y-axes in Figure 6). When >50% of coauthor pairs comprise one early-career and one established researcher, we expect gender heterophily (*α* < 0) whenever the gender ratio differs between career stages. Conversely, when >50% of collaborations are between people at the same career stage, we expect gender homophily (*α* > 0). In a few parameter spaces (shown in red; Figure 6), *α* was quite high, and overlapped with the values that we estimated (Figure 2).

**Figure 6:**
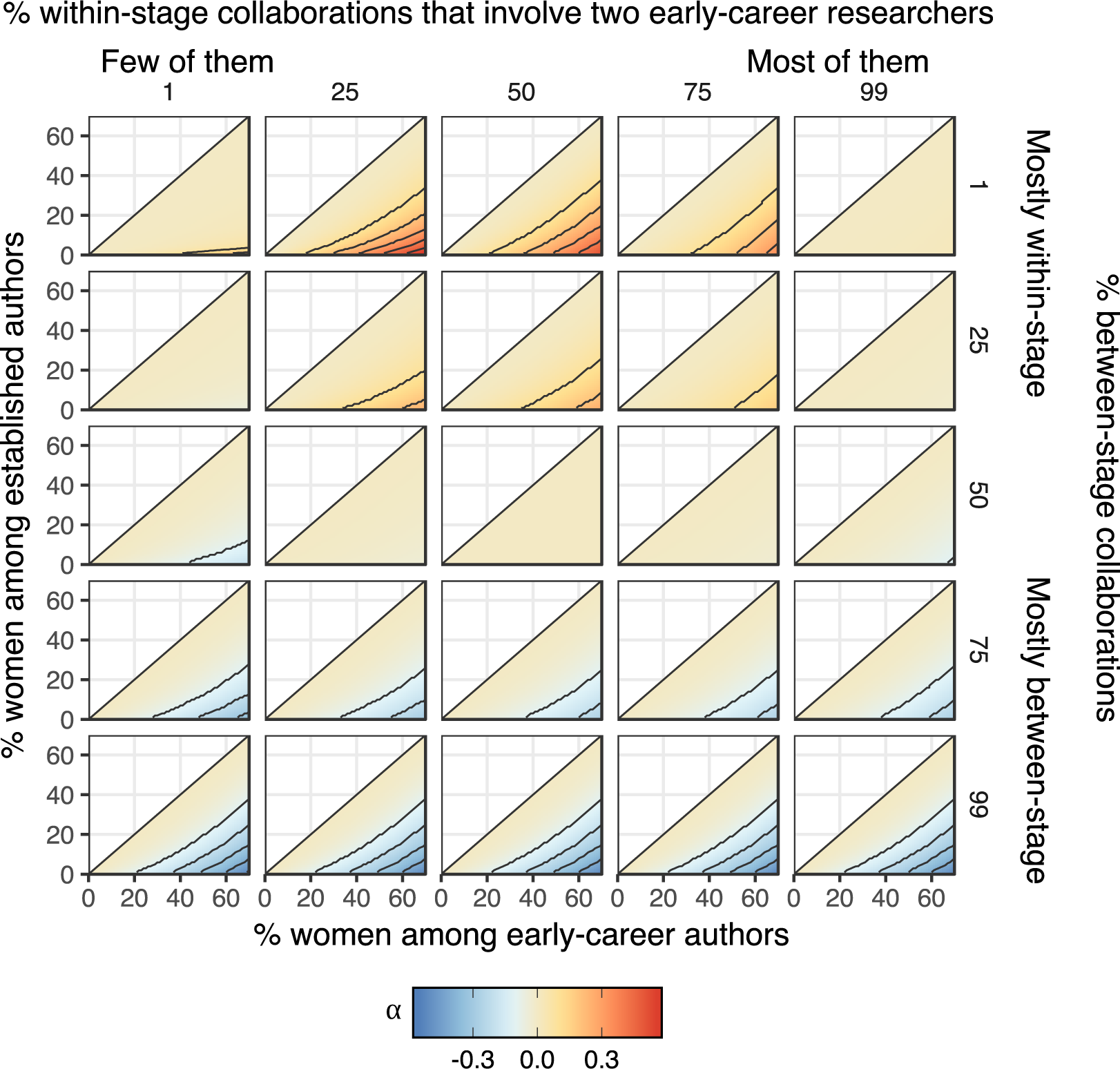
When the gender ratio of early-career researchers is not equal to the gender ratio among established researchers, the null expectation for *α* is not necessarily zero. Specifically, if most collaborations occur between career stages, there will be an excess of mixed-gender collaborations (*α* < 0, blue areas), while if most collaborator pairs comprise two people at the same career stage, there will be an excess of same-gender collaborations (*α* > 0, red areas). However, the conditions required for strong gender homophily (i.e. the red areas) are quite restrictive, making it unlikely that this issue can fully explain the homophily observed in our study. Additionally, in research disciplines where between-career stage collaboration is common and there is a shortage of women among established researchers (i.e. the blue areas), our study will underestimate the strength of gender homophily. Contour lines denote increments of 0.1.

Despite this overlap, Figure 6 suggests that our main conclusions (and those of other studies of gender homophily) are probably robust to this career stage issue. We only expect strongly positive *α* when A) the gender ratio is highly skewed across career stages (e.g. a 5-fold difference), and B) collaborations between early and established researchers are very rare (e.g. <10% of the total). Both of these conditions seem unlikely to be true for most fields: the gender gap across careers stages is generally less pronounced [1,5], and it is very common for early-career researchers to co-publish with an established mentor [61]. However, one can get *α* > 0 for realistic combinations of parameters, e.g. a moderate shortage of women in senior positions coupled with a moderate excess of within-career stage collaboration, suggesting this effect might contribute to some of the homophily observed by this and previous studies.

Lastly, we note that if there is a gender gap between career stages and coauthorships between early-career and established researchers comprise >50% of the total, then the baseline expectation for *α* is actually less than zero (blue areas in Figure 6). Therefore, it is possible that researchers preferentially assort with same-gendered collaborators even more strongly than implied by our results, at least for certain journals or research disciplines.

## Discussion

We found evidence that researchers work with same-gendered coauthors more often than expected under the null model, even after implementing stringent controls for Wahlund effects (Figure 1). Our study therefore reaffirms earlier studies’ conclusions [49–57,62] using stricter methodology, and generalises their results across the life sciences. Relatively few journals had *α′* values below zero, and almost no journals showed statistically significant gender heterophily after controlling for multiple testing. The excess of same-gender coauthorships was quite large: many journals had *α′* > 0.1, indicating that the gender ratio of men’s and women’s coauthors differs by >10% in absolute terms. In relative terms, our findings are even more striking: for example, if men have 20% female coauthors and women have 30% (i.e. *α′* = 0.1 in a field with a typical gender ratio [5]), then women publish with women 50% more often than men do.

An important limitation of our study is that we cannot reliably determine the cause(s) of the observed excess of same-gender coauthorships. As well as the obvious interpretation – conscious or unconscious selection of same-gender collaborators by men, by women, or both – our results could be partly explained by uncontrolled Wahlund effects. However, we suspect the contribution of these uncontrolled artefacts to be minor, for four reasons: we found positive *α′* after controlling for three obvious sources of Wahlund effect; there was no inflation of *α′* in highly multidisciplinary journals relative to specialised journals; restricting the data by country yielded similar estimates of *α′*; and we used a simple model to show that differences in gender ratio between career stages are unlikely to fully explain our results. On balance, we believe the data suggest that it is likely that some researchers preferentially select same-gendered collaborators, although it is difficult to ascertain what proportion of people show such a preference, or how much the strength of the preference varies between individuals. We also note that even in a world in which collaboration was completely random with respect to gender, a high proportion of individual researchers would have entirely same-gendered collaborators by chance alone (especially in gender-biased disciplines); thus, individuals who only have same-gendered co-authors are not necessarily doing anything differently from people with gender-balanced co-authors.

We hypothesised that disciplines with a strongly skewed gender ratio might show the strongest gender homophily, e.g. because being in the minority might increase people’s motivation to seek out same-gendered colleagues. Contrary to this hypothesis, we found no evidence that gender homophily is restricted to particular disciplines: *α′* was similarly high across the board (Figure 2). Interestingly, gender homophily was weakest for journals with a male-biased author gender ratio, and strongest in journals with a female-biased author gender ratio. One possible reason is that men are more likely to preferentially seek out male collaborators in fields where men are a minority, relative to the homophily displayed by women in fields where women are a minority. However, this latter result only has tentative statistical support since our sample contains few journals in which most authors are women (Figure 4).

We also found that gender homophily was marginally stronger in 2015-2016 relative to 2005-2006. Although this trend might reflect a change in the gender preferences of researchers seeking collaborators, there are alternative (and perhaps more likely) explanations. For example, this trend might result from the increasing number of women working in senior positions in STEMM over the past decade [63–65]. As shown in Figure 6, if enough coauthorships are between junior and senior researchers, a large gender gap between career stages can give the appearance of heterophily. As this gender gap between career stages lessens, the observed values of *α′* may increase.

Regarding our finding of weaker homophily among 2-author papers, we suspect that many 2-author teams comprise a student/postdoc and a senior staff member, making these teams especially likely to be mixed-gender, due to the greater shortage of women among senior researchers [1,5]. Assuming this interpretation is correct, this result suggests that our reported *α′* values may underestimate the strength of peoples’ preferences for same-gendered collaborators; essentially, women seeking a senior collaborator could be constrained to work mostly with men, meaning that people’s ideal and realised gender preferences would be mismatched. On a related note, Ghiasi et al. [51] argue that women in engineering are “compliant [in reproducing] male-dominated scientific structures” because they do not collaborate often enough with other women (for reference, Figure 7 in [51] implies that coauthorships involving two women are *c*. 30% more frequent than expected under random assortment). By contrast, we feel that it may be counter-productive to recommend that women collaborate primarily with other women, e.g. because this constrains women’s options (particularly in fields like engineering, where 90% of professors are men [1]). Instead, we suggest that researchers of both genders can help to close the gender gap in STEMM. In the context of collaboration, one way to do this is to undertake self-examination to ensure that one is not inadvertently overlooking or excluding women among potential students and colleagues. One should also take care to treat male and female collaborators equally, e.g. in terms of training and mentoring, allocation of work, how one frames or promotes the collaboration (e.g. in conference presentations or on a website), and in terms of the credit afforded to collaborators’ contributions. Experimental work suggests that unconscious bias causes people to undervalue women’s research achievements [20], and a study of author contribution statements found observational evidence that menial or under-valued tasks are assigned to women while more prestigious tasks are assigned to men [61].

Our study begs two questions: what causes gender homophily in science, and are our results cause for concern? We believe that the answers to these questions are closely related. For example, some of the homophily we observed might be caused by women seeking to avoid harassment or sexism from men [38], which would clearly be concerning. Additionally, Sheltzer and Smith [66] concluded that ‘elite’ male academics (defined as recipients of major honours) have a higher proportion of male students and postdocs than non-elite male academics. This finding could contribute to the homophily we observed, and is cause for concern since the results might reflect discrimination against women during hiring [20], or avoidance by women of elite research groups (e.g. due to gender differences in confidence, or a perception that some groups are sexist). We also found a little evidence that gender homophily is detrimental to research quality, in that high-impact journals tended to have weaker homophily. Assuming that papers published in high-impact journals are of higher average quality (which is contentious; [67]), our results provide non-experimental support for the hypothesis that mixed-gender teams produce better research than single-gender teams [42–48]. Another issue is that if many collaborations are between established researchers, there will be an excess of male-male collaborations in fields where women in senior positions are rare; some of the observed homophily might therefore reflect the elevated gender gap among senior researchers.

On the other hand, homophily might have more benign causes. Collaboration is often most enjoyable and productive when working with like-minded people, who might tend to be same-gendered more often than not. We also suppose that some people consciously choose to preferentially collaborate with women in order to help close the gender gap in the workforce; this would create homophily if women do this more than men. In support of this interpretation, there is some evidence that women are more likely than men to promote the work of female colleagues by inviting them to give talks [68,69]. Given that many collaborative research projects unfortunately involve a gendered division of labour [61], working with a same-gendered colleague may provide exposure to new parts of the research process, and (especially for the minority gender) a welcome change of pace.

## Methods

### The dataset

We used the dataset of PubMed author lists from Holman et al. [5]. Briefly, that dataset was created by downloading every article indexed on PubMed and attempting to infer each author’s gender from their given name using computational methods. Each journal was assigned to one of 107 scientific disciplines, using PubMed’s journal categorisations in the interests of objectivity. Because the present study focuses on co-authorship, all single-author papers were discarded. We also discarded all papers for which we could not determine the gender of every author with *≥*95% certainty, in order to simplify the statistical analysis. To mitigate Wahlund effects caused by variation in the gender ratio of researchers over time (see below), we only kept papers with publication dates falling in two one-year time periods, namely 0-1 or 10-11 years prior to the collection date of the PubMed data (i.e. 20^th^ August 2016). Lastly, we excluded journals with fewer than 50 suitable papers. Detailed sample size information is given in S1 Table.

### Calculating *α*, the coefficient of homophily

Following Bergstrom et al. [60], we defined the coefficient of homophily as *α = p − q*, where *p* is the probability that a randomly-chosen co-author of a *male* author is a man and *q* is the probability that a randomly-chosen co-author of a *female* author is a man. Like the Wahlund effect, *α* is borrowed from population genetics; for a set of 2-author papers, it is equivalent to Wright’s coefficient of inbreeding [70]. Mathematical work illustrates that *α* is closely related to alternative network-based methods for quantifying homophily [71].

To estimate *α* for a particular subset of the scientific literature, we estimated *p* as the average proportion of men’s co-authors who are men (averaged across all papers with at least one man author), and *q* as the average proportion of women’s co-authors who are men (averaged across all papers with at least one woman author). To estimate the 95% confidence intervals on *α* for a given set of *n* papers, we sampled *n* papers with replacement 1000 times, estimated *α* on each sample, and recorded the 95% quantiles of the resulting 1000 estimates.

As well as calculating *α* for all authors, we calculated *α* for first or last authors only. *α* was again defined as *p − q*, but this time *p* was estimated as the average proportion of male co-authors on papers with a male first (or last) author, and *q* was estimated as the average proportion of male co-authors on papers with female first (or last) authors. We did not calculate *α* for other authorship positions (e.g. second or third authors) because this would necessitate culling the dataset to include only papers with a sufficiently long author list, complicating interpretation of the results.

We also calculated *α* for papers with 2, 3, 4 or *≥*5 authors, for all journals that had at least 50 suitable papers from 2015-2016 with the specified author list length.

Our test assumes that the expected value of *α* is zero if authors randomly assort, but for small datasets this assumption is not always true. Essentially, this issue arises because a person cannot be their own co-author. In a small dataset comprising *m* men and *f* women, a man can co-author with *m* − 1 men while women can co-author with *m* men. Thus, the null expectation for *alpha* is a negative number – potentially a large one if *m* and *f* are very small.

To control for the fact that the null expectation for *α* is not zero for small datasets, we devised an adjusted version of the coefficient of homophily, which we term *α′*. Every time we calculated *α* for a set of papers, we also determined the expected value of *α* under the null hypothesis that authors assort randomly with respect to gender. This was accomplished by randomly permuting authors across papers 1000 times, recalculating *α*, and taking the median. We then calculated *α′* by subtracting the null expectation for *α* from the observed value. We also used the null-simulated *α* values to calculate a two-tailed p-value for the observed value of *α*; the p-value was defined as the proportion of null simulations for which |*α_null_*| > |*α_obs_*|. We applied false discovery rate (FDR) correction to each set of p-values to account for multiple testing [72].

As expected, *α′* was usually almost identical to *α* (S7 Fig), but *α* was downwardly biased relative to *α′* for small datasets (S8 Fig). Additionally, the correlation between *α′* and sample size was negligible (*R*^2^ < 0.01), suggesting that our calculation of *α′* effectively removed the dependence of *α* on sample size. We therefore used *α′* in all analyses.

### Minimising the Wahlund effect: research discipline and time period

To minimise bias in *α′* due to the Wahlund effect, we restricted each set of papers to a single research specialty to the greatest extent allowed by our data. Specifically, we only calculated *α′* for individual journals, since papers from the same journal typically focus on closely related topics. Although some journals, e.g. *PLOS ONE*, publish research from diverse disciplines with very different author gender ratios [5], calculating *α′* for these highly multidisciplinary journals is still useful as a contrast. The difference in *α′* between highly multidisciplinary and more specialised journals, e.g. *PLOS ONE* versus *PLOS Computational Biology*, gives an estimate of the extent to which multidisciplinarity within journals inflates *α′*.

As well as varying between disciplines, the gender ratio of authors has changed markedly over time [5]. Because the gender ratio was more male-biased in the past, *α′* would be inflated if we calculated it for a sample of papers published over a long enough time frame. To minimise this effect, we only sampled papers from two one-year periods (namely 2005-6 and 2015-16). The median change per year in % (fe)male authors across journals is below 0.5% [5], and so restricting our dataset to a single year should prevent temporal changes in gender ratio from noticeably affecting our estimates of *α′*.

### Minimising the Wahlund effect: author country of affiliation

A Wahlund effect could arise even if one calculates *α′* for a single discipline and time period, because of variation in the gender ratio of researchers from different countries. For example, Holman et al. [5] found that PubMed-indexed authors based in Serbia are more than twice as likely to be women as are authors based in Japan. Therefore, a dataset containing a mix of papers from teams of authors based in these two countries would contain an excess of same-sex coauthorships, even if collaboration were random with respect to gender within each country.

To address this issue, we also analysed every combination of journal and author country of affiliation for which we had enough data (i.e. 50 or more papers published in 2015-16). For simplicity, we restricted the dataset to only include papers for which Holman et al. [5] had identified the country of affiliation for all authors on the paper, and all authors shared the same country of affiliation. Restricting the dataset in this fashion produced enough data to measure *α′* for 325 combinations of journal and country (median: 70 papers and 273 authors per combination).

### Calculating standardised journal impact factor

We obtained the 3-year impact factor for each journal from Clarivate Analytics (formerly ISI). To account for large differences in impact factor between disciplines, we took the the residuals from a model with *log*_10_ impact factor as the response and the research discipline of the journal as a random effect. Thus, journals with a positive standardised impact factor have a higher mean number of citations than the average for journals in their discipline. We then used Spearman rank correlation to test whether *α′* was correlated with impact factor across journals.

### Statistical analysis

Previous authors [66,73] have hypothesised that senior scientists preferentially recruit staff and students of the same gender, and/or that junior researchers preferentially select same-gendered mentors. In the majority of PubMed-indexed disciplines, authorship conventions mean that the first-listed author is often an early-career researcher, while the author listed last is more likely to be a senior researcher leading a research team [74]. Assuming that senior researchers are the main drivers of homophily and that there are enough papers with three or more authors, we predict that the last author’s gender will be the strongest predictor of the remaining authors’ genders (i.e. the gender of the last author will be more salient than that of the first author, or any other authorship position). This is because the first author’s gender would simply be an imperfect correlate of the true causal effect, while the last author’s gender would be the causal effect itself.

To test whether *α′* for last authors tends to be higher than *α′* for first authors for any given dataset, we used a linear mixed model implemented in the lme4 and lmerTest packages for R, with *authorship position* (first or last) as a fixed factor, and *journal* and *research discipline* as crossed random effects. The response variable was *α′*, and we weighted each observation by the inverse of the standard error from our estimate of *α′*, meaning that more accurate measurements of *α′* had more influence on the results. We used a similar model to test for a difference in *α′* between the 2005-6 and the 2015-16 datasets, with two differences: we fit year range as a two-level fixed factor (instead of authorship position), and we used *α′* estimated for all authors (not first/last authors) as the response variable.

The relationship between the gender ratio of authors publishing in a journal and its *α′* value appeared nonlinear (see Results). We therefore fit a generalised additive model with thin plate regression spline smoothing, implemented using the mgcv package for R.

To model the relationship between *α′* and the number of authors on the paper, we used a meta-regression model implemented in the R package brms [75]. The model incorporated the standard error associated with each estimate of *α′*, had author number as a fixed effect, and journal as a random intercept (to control for repeated measures of each journal). We also fit a random slope of author number within journal, thereby allowing the response to author number to vary between journals. We used the default (weak) priors. The full output of this model can be viewed in the Online Supplementary Material.

### Theoretical expectations for *α* when the gender ratio differs between career stages

In most STEMM disciplines, the gender ratio is more skewed among established researchers relative to early-career researchers, due both to women leaving STEMM careers at greater rates (the ‘leaky pipeline’), and to historical shortages of women studying STEMM subjects at university (‘demographic inertia’) [1,5]. We hypothesised that this difference in gender ratio between career stages could potentially create both Wahlund effects and ‘reverse’ Wahlund effects. For example, imagine that the majority of collaborations in a particular field are between students and professors, and that the gender ratio differs between career stages: we would then see an excess of mixed-gender coauthorships (heterophily, *α* < 0), even if gender has no direct, causal effect. Similarly, a hypothetical field in which students work only with students, and professors with professors, would have apparent gender homophily (*α* > 0).

We can think of no tractable method of controlling for this issue using our dataset, which contains no information on career stage. Therefore, we instead decided to derive theoretical expectations for *α* when there is a difference in gender ratio across career stages, in order to determine if and how this effect should alter our inferences. For simplicity, our calculations assume there are only two career stages (‘early-career’ and ‘established’), though we expect that the general conclusions would also apply to a multi-tier career ladder. Under the null model that gender has no causal effect on collaboration, we calculated *α* for various combinations of the four free parameters in our simple model. These parameters are: the gender ratio among early-career researchers (x-axis of Figure 6), the gender ratio among established researchers (y-axis of Figure 6), the frequency of within-versus between career stage collaborator pairs (rows in Figure 6), and lastly the frequency of within-stage collaborations that are between two early-career researchers as opposed to two late-career researchers (columns in Figure 6). When these four parameters are specified, one can easily calculate the relative frequencies of collaborator pairs that involve two men, two women, or a man and a woman. In short, if we have specified the frequency of women at both career stages, as well as the frequency of the three possible types of collaboration with respect to career stage (early-early, early-established, and established-established), then we can calculate the frequency of collaborators pairs comprising two women, or a woman and a man, and thus find *α* (see the Online Supplementary Material for the annotated R code).

## Data availability and reproducibility

The Online Supplementary Material contains R scripts used to produce all results, figures and tables; it can be viewed online at https://lukeholman.github.io/genderHomophily/. The input data from Holman et al. [5] is archived at https://osf.io/bt9ya/.

## Acknowledgements

CM was supported by the Academy of Finland (284666 to the Centre of Excellence in Biological Interactions). We thank Devi Stuart-Fox and Dominique Potvin for helpful discussion.

## Supporting information

### Supplementary figures

S1 Fig. Plot showing the percentage of papers that have 1, 2, 3, 4, or *≥*5 authors for each discipline in the dataset of Holman et al. (2018). This information can also be found in S3 Data.

S2 Fig. Histogram showing the distribution of differences in *α′* between the 2015-16 and 2005-6 samples, where positive numbers indicate an increase in *α′* with time. The mean is slightly positive (i.e. 0.004), indicating a mild increase in average *α′* with time.

S3 Fig. Histogram showing the difference between *α′* calculated for first and last authors. Positive values mean that *α′* was higher when calculated for first authors, and negative values mean *α′* was higher when calculated for last authors. The mean is very slightly higher than zero, indicating that *α′* tends to be higher for first authors.

S4 Fig. Histogram of *α′* for 325 unique combinations of journal and country, using data from August 2015 - August 2016. The white areas denote combinations for which *α′* differs significantly from zero (p < 0.05, following false discovery rate correction).

S5 Fig. Plot showing the 68 combinations of journal and author country of affiliation for which *α′* is significantly higher than expected.

S6 Fig. Histogram showing the estimated degree to which *α′* is inflated by inter-country differences in author gender ratio, across the 285 journals for which we had adequate data after restricting the analysis by country. The average inflation in *α′* is negligible, suggesting that Wahlund effects resulting from inter-country differences have a negligible effect on our estimates of gender homophily.

S7 Fig. There is a very strong correlation between the values of *α* and *α′* calculated for each journal, though in a handful of cases the difference is considerable. The deviation between *α* and *α′* is greatest for journals for which there is a small sample size (see S8 Fig).

S8 Fig. For journals for which we recovered a small number of papers (<100), the unadjusted metric *α* was downwardly biased. This fits our expectations: because researchers cannot be their own co-authors, small datasets will tend to produce negative estimates of *α* even if authors assort randomly with respect to gender (see main text). This suggests that *α′* is a better measure of homophily and heterophily, though the improvement is trivial in large enough samples.

### Supplementary tables

S1 Table. Sample sizes for the two datasets, which comprise papers published in the timeframes August 2005 - August 2006, and August 2015 - August 2016.

S2 Table. Number of journals showing statistically significant homophily or heterophily, in two one-year periods. The significance threshold was p < 0.05, and p-values were adjusted using Benjamini-Hochberg false discovery rate correction. Note that the power of our test is lower for the 2005-2006 data because fewer papers were recovered per journal: thus, it is not meaningful to compare the % significant journals (i.e. 11% vs 24%) between the two time periods.

### Supplementary datasets

S1 Data: This spreadsheet shows the *α* values calculated for each journal, in the 2005 and 2015 samples, and for each type of author (all authors, first authors, and last authors). The tables gives the impact factor of each journal, the sample size, *α* and *α′* and their 95% CIs, and the p-value from a 2-tailed test evaluating the null hypothesis that *α* is zero (both raw and FDR-corrected p-values are shown).

S2 Data: This file gives the number and percentage of papers that have 1, 2, 3, 4, or *≥*5 authors for each *journal* in the dataset of Holman et al. (2018) *PLOS Biology*. Note that the sample sizes include papers for which the gender of one or more authors was not determined by Holman et al.

S3 Data: This file gives the number and percentage of papers that have 1, 2, 3, 4, or *≥*5 authors for each *discipline* in the dataset of Holman et al. (2018) *PLOS Biology*. Note that the sample sizes include papers for which the gender of one or more authors was not determined by Holman et al.

S4 Data. The table shows the distribution of the *α′* values across journals, split by the research discipline. The gender ratio column shows the percentage of women authors in the sample used to calculate *α′*, across all authorship positions. In the last two columns, the numbers outside parentheses give the number of journals that deviate statistically significantly from zero, while the numbers inside parentheses give the number that remain significant after false discovery rate correction.

